## Supplemental tables, figures and methods for "Novel artificial selection method improves function of simulated microbial communities"

### Supplementary tables

| Method 1 | Method 2 | $p$ (IBM) | $p$ (ODE) |
|---|---|---|---|
| DS | $\emptyset$ | $4\times10^{-10}$ | $4\times10^{-10}$ |
| DS | PS | $8\times10^{-10}$ | $8\times10^{-10}$ |
| DS | MS | $8\times10^{-10}$ | - |
| DS | PIS | $1\times10^{-9}$ | $2\times10^{-8}$ |
| DS | MIS | $8\times10^{-10}$ | - |
| DS | DR | $8\times10^{-10}$ | $8\times10^{-10}$ |
| DS | NS | $8\times10^{-10}$ | $8\times10^{-10}$ |
| PS | $\emptyset$ | 1.0 | 1.0 |
| PS | MS | $1\times10^{-7}$ | - |
| PS | PIS | $4\times10^{-9}$ | $7\times10^{-5}$ |
| PS | PR | $8\times10^{-10}$ | $3\times10^{-9}$ |
| PS | NS | $8\times10^{-10}$ | 0.1 |
| MS | $\emptyset$ | 1.0 | - |
| MS | MIS | 0.67 | - |
| MS | MR | $8\times10^{-10}$ | - |
| MS | NS | 0.3 | - |
| PIS | $\emptyset$ | $9\times10^{-6}$ | 0.8 |
| MIS | $\emptyset$ | 1.0 | - |

**Table S1.** P-values for Fig.2A-D a Wilcoxon signed-rank test of difference in maximum degradation between methods.

| Method 1 | Method 2 | $p$ (IBM) | $p$ (ODE) |
|---|---|---|---|
| DS | PS | $8\times10^{-10}$ | $7\times10^{-10}$ |
| DS | MS | $8\times10^{-10}$ | - |
| DS | PIS | $8\times10^{-10}$ | $7\times10^{-10}$ |
| DS | MIS | $8\times10^{-10}$ | - |
| DS | DR | $8\times10^{-10}$ | $8\times10^{-10}$ |
| DS | NS | $8\times10^{-10}$ | $7\times10^{-10}$ |
| PS | MS | $1\times10^{-4}$ | - |
| PS | PIS | $3\times10^{-8}$ | $7\times10^{-4}$ |
| PS | PR | $2\times10^{-9}$ | $7\times10^{-7}$ |
| PS | NS | $8\times10^{-10}$ | $3\times10^{-6}$ |
| MS | MIS | $8\times10^{-2}$ | - |
| MS | MR | $2\times10^{-8}$ | - |
| MS | NS | $2\times10^{-9}$ | |

**Table S2.** P-values for Fig.2E-H a Wilcoxon signed-rank test of difference in degradation ranks between methods.

| Method 1 | Method 2 | $p$ (IBM) | $p$ (ODE) |
|---|---|---|---|
| DS | PS | $6 \times 10^{-9}$ | $8 \times 10^{-10}$ |
| DS | MS | $8 \times 10^{-10}$ | - |
| DS | PIS | $8 \times 10^{-10}$ | $8 \times 10^{-10}$ |
| DS | MIS | $8 \times 10^{-10}$ | - |
| DS | DR | $8 \times 10^{-10}$ | $6 \times 10^{-4}$ |
| DS | NS | $8 \times 10^{-10}$ | $8 \times 10^{-10}$ |
| PS | MS | $1 \times 10^{-9}$ | - |
| PS | PIS | 0.1 | 0.01 |
| PS | PR | $2 \times 10^{-8}$ | 0.5 |
| PS | NS | $8 \times 10^{-10}$ | 0.3 |
| MS | MIS | 0.13 | - |
| MS | MR | $1 \times 10^{-9}$ | - |
| MS | NS | $1 \times 10^{-5}$ | - |

**Table S3.** P-values from a Wilcoxon signed-rank test of difference in total investment in communities, between methods for Fig.3A-D.

### Supplementary figures

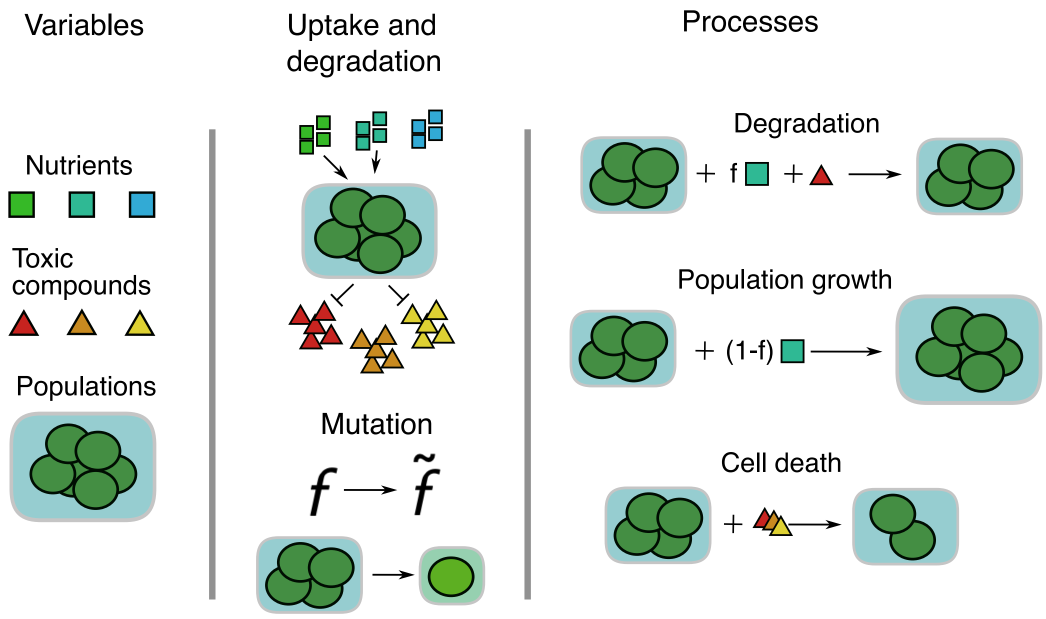


**Fig. S1.** Illustration of the variables and processes in the ODE model. Populations of cells vary in their preferences for nutrients and degradation capabilities. The populations use the available nutrients to degrade the toxic compounds and to grow.

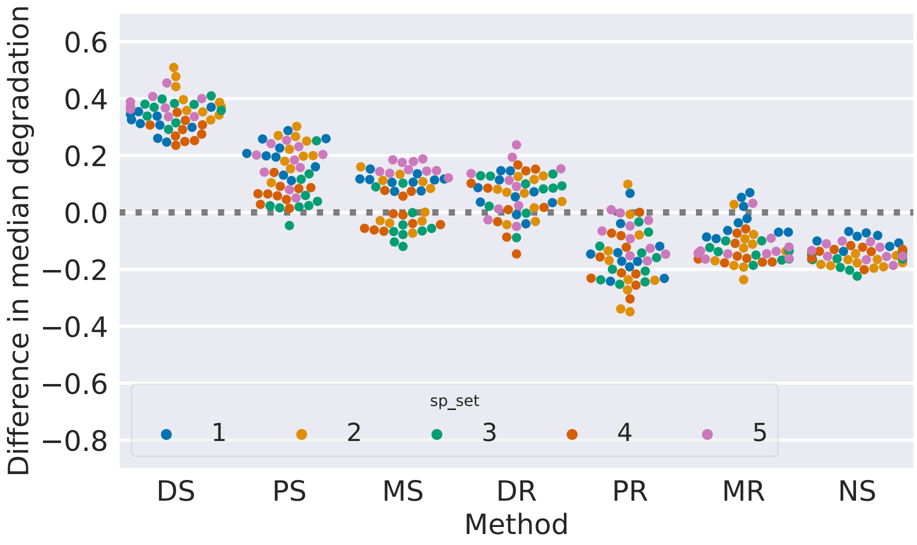


**Fig. S2.** The difference in **median** degradation between round 50 and round 0 for each propagation method, corresponding to Fig. 2 where the difference in maximum degradation is shown. The two-sided Wilcoxon test for difference in degradation against the no-selection control is significant for the selection methods DS, PS and MS. Data generated by the IBM.

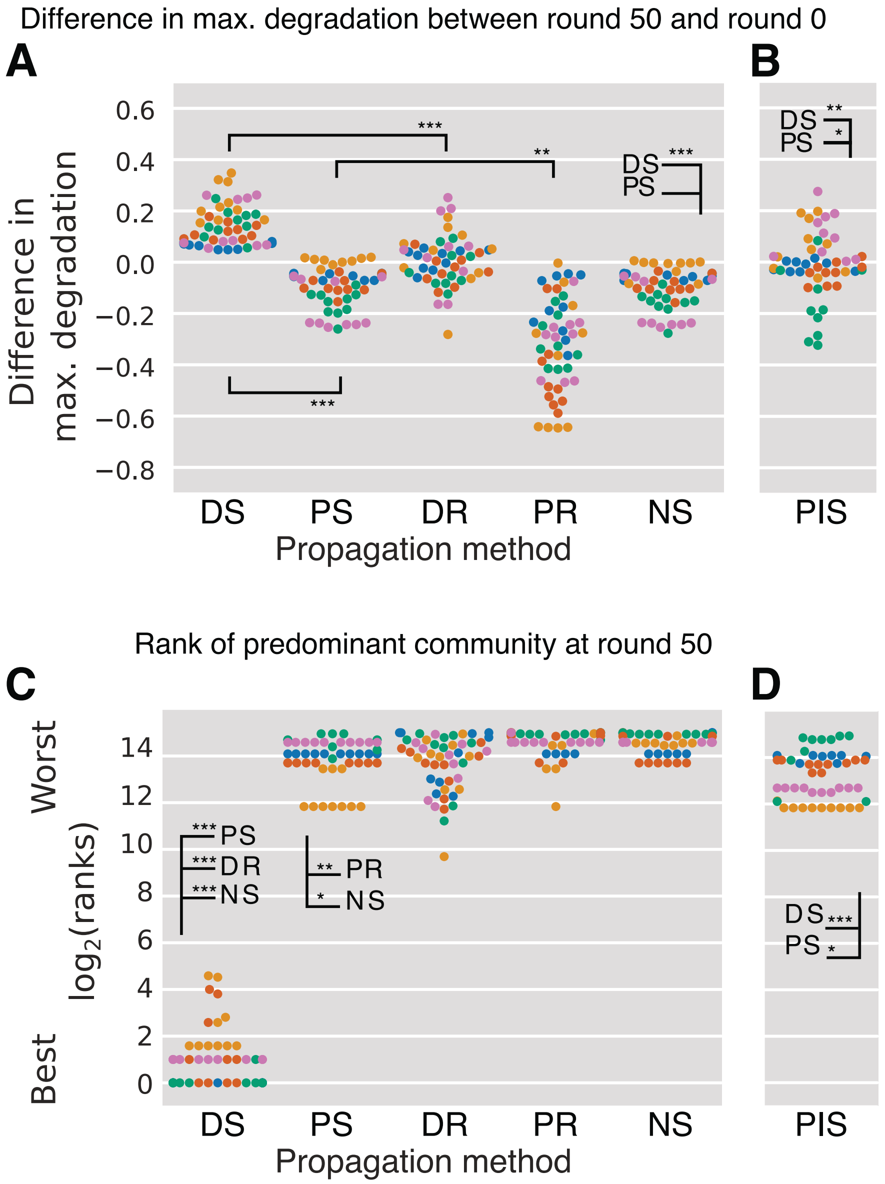


**Fig. S3.** Degradation scores and ranking of selected communities. (A, B) The difference in maximum degradation between round 50 and round 0 for each propagation method, corresponding to Fig. 2, but generated by the ODE. (C, D) The rank of the predominant community (the most common combination of species among the 21 communities in the last round of selection, not counting sub-communities) in terms of its degradation score compared to all of the 32767 possible combinations of 1, 2, ..., 15 ancestral species, corresponding to Fig. 2, but generated by the ODE.

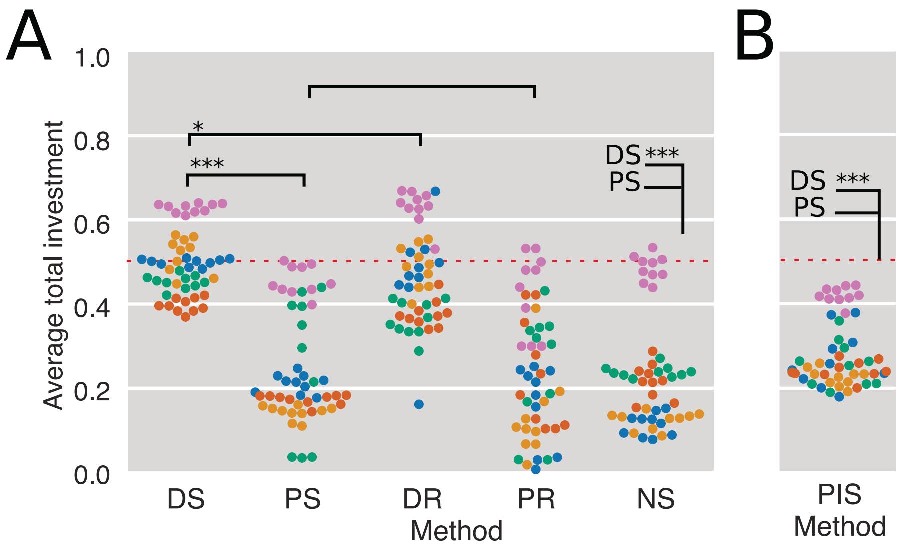


**Fig. S4.** Average total investment into degradation of communities at round 50, averaged for the 21 communities in a run, corresponding to panel A, B in Fig. 3, but generated by the ODE.

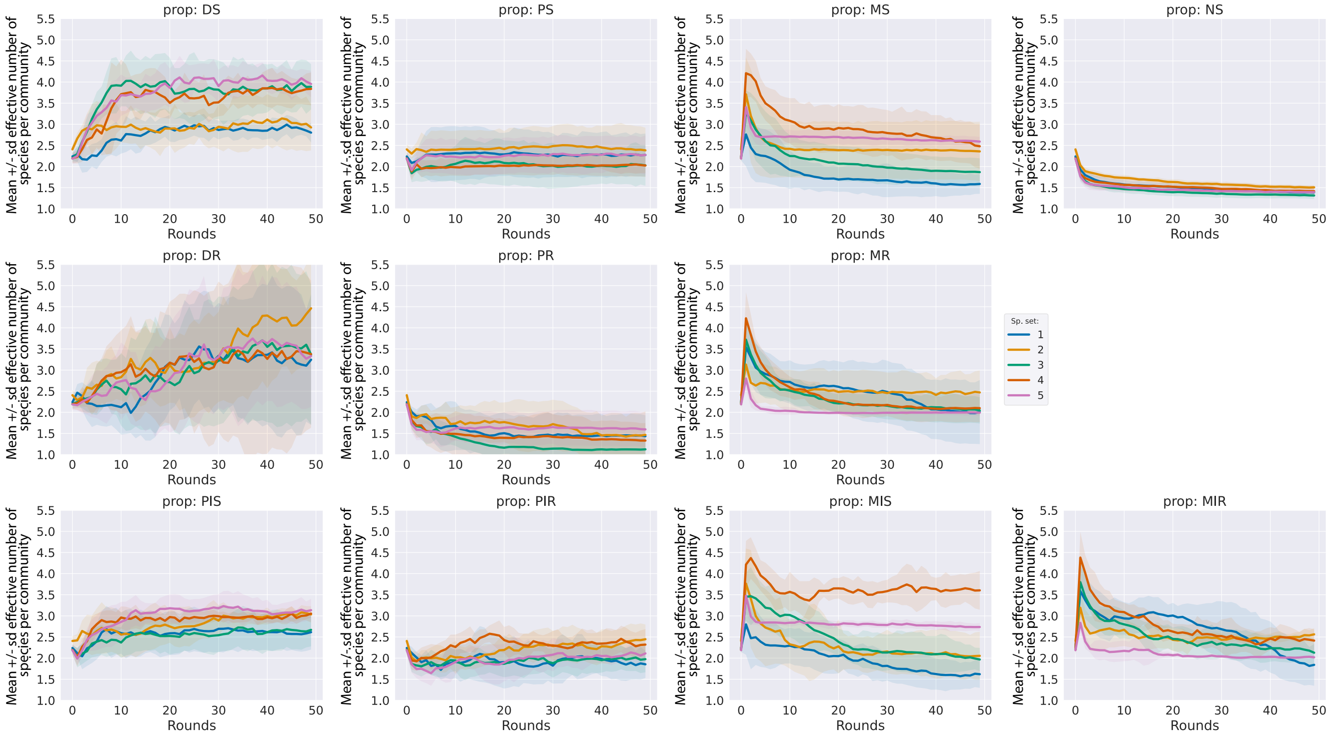


**Fig. S5.** Time-series of the species diversity (effective number of species per community) corresponding to Fig. 3C. Each panel shows the mean $\pm$ standard deviation over the 10 repeated runs, for each species set 1-5, for one propagation method. Data generated by the IBM.

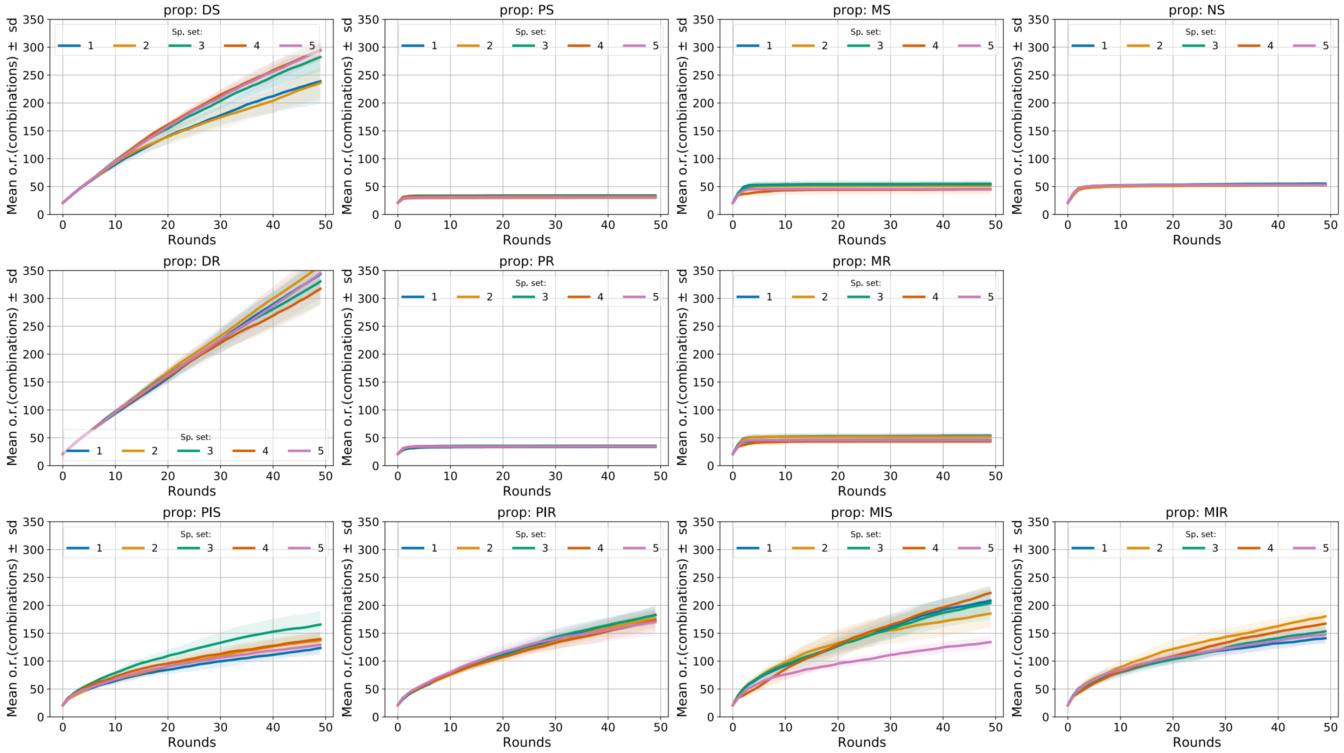

**Fig. S6.** Time-series of the number of explored communities, corresponding to Fig. 4A. Each panel shows the mean $\pm$ standard deviation over the 10 repeated runs, for each species set 1-5, for one propagation method. Data generated by the IBM.

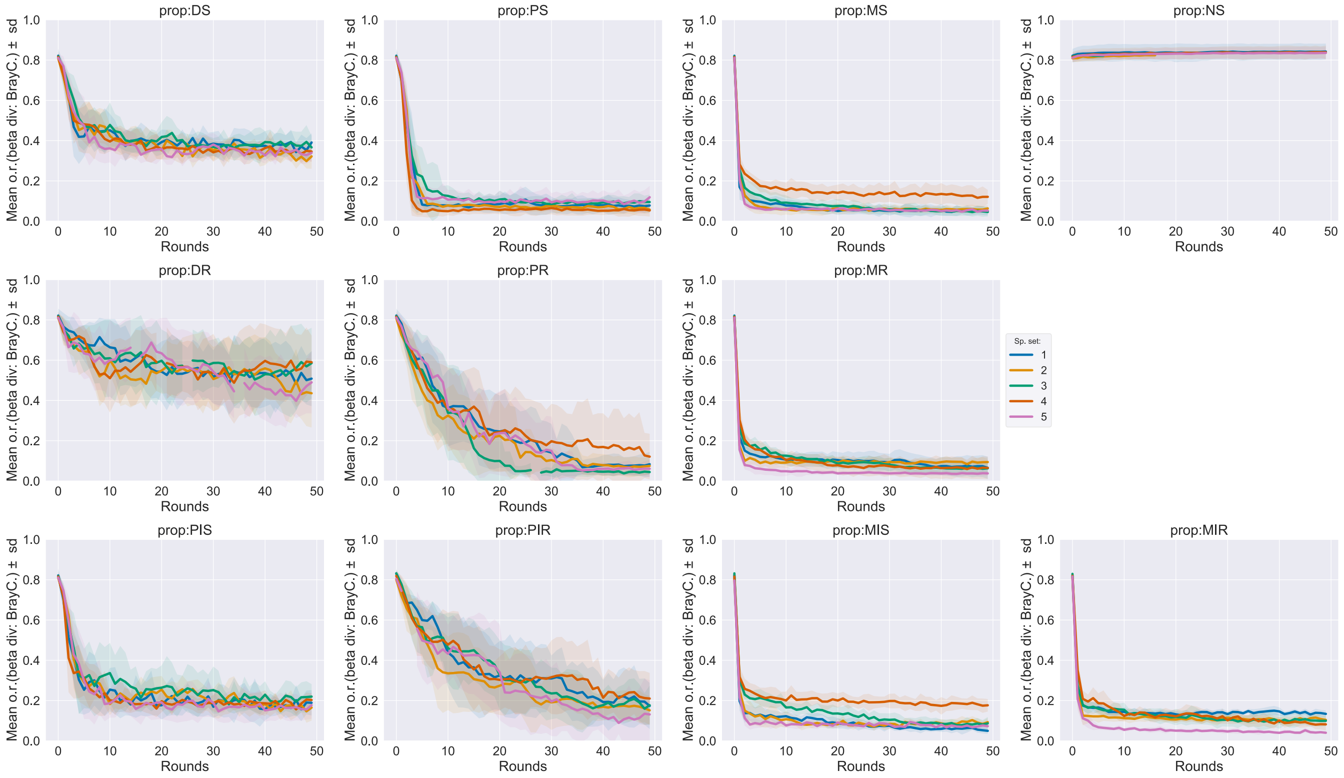

**Fig. S7.** Time-series of the beta diversity corresponding to Fig. 4B. Each panel shows the mean $\pm$ standard deviation over the 10 repeated runs, for each species set 1-5, for one propagation method. Data generated by the IBM.

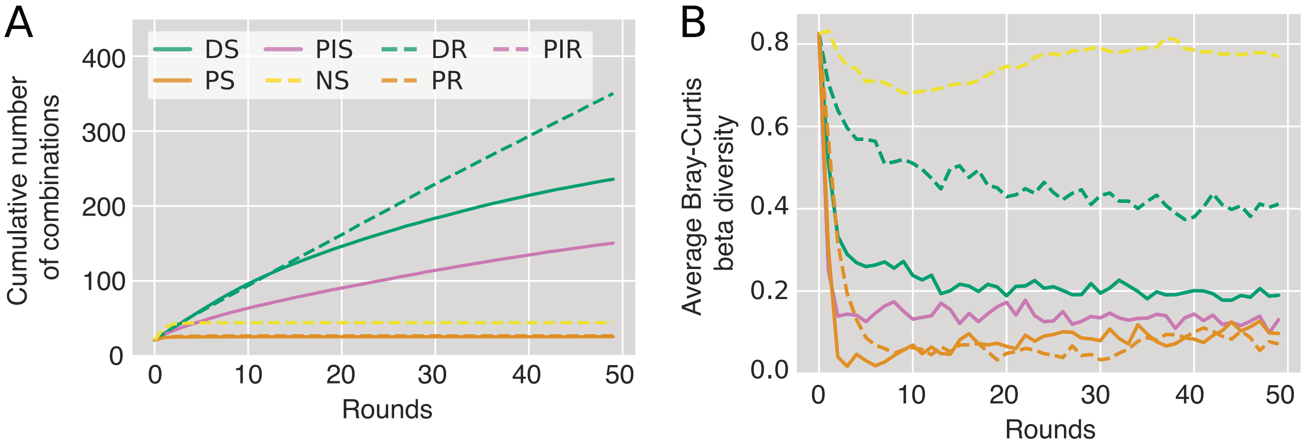


**Fig. S8.** (A) Time-series of cumulative number of unique communities for each selection method, corresponding to Fig. 4A but generated using the ODE model. Time-series of the beta diversity corresponding to Fig. 4B, but generated using the ODE model. Each panel shows the mean ± standard deviation over the 10 repeated runs, for each species set 1-5, for one propagation method.

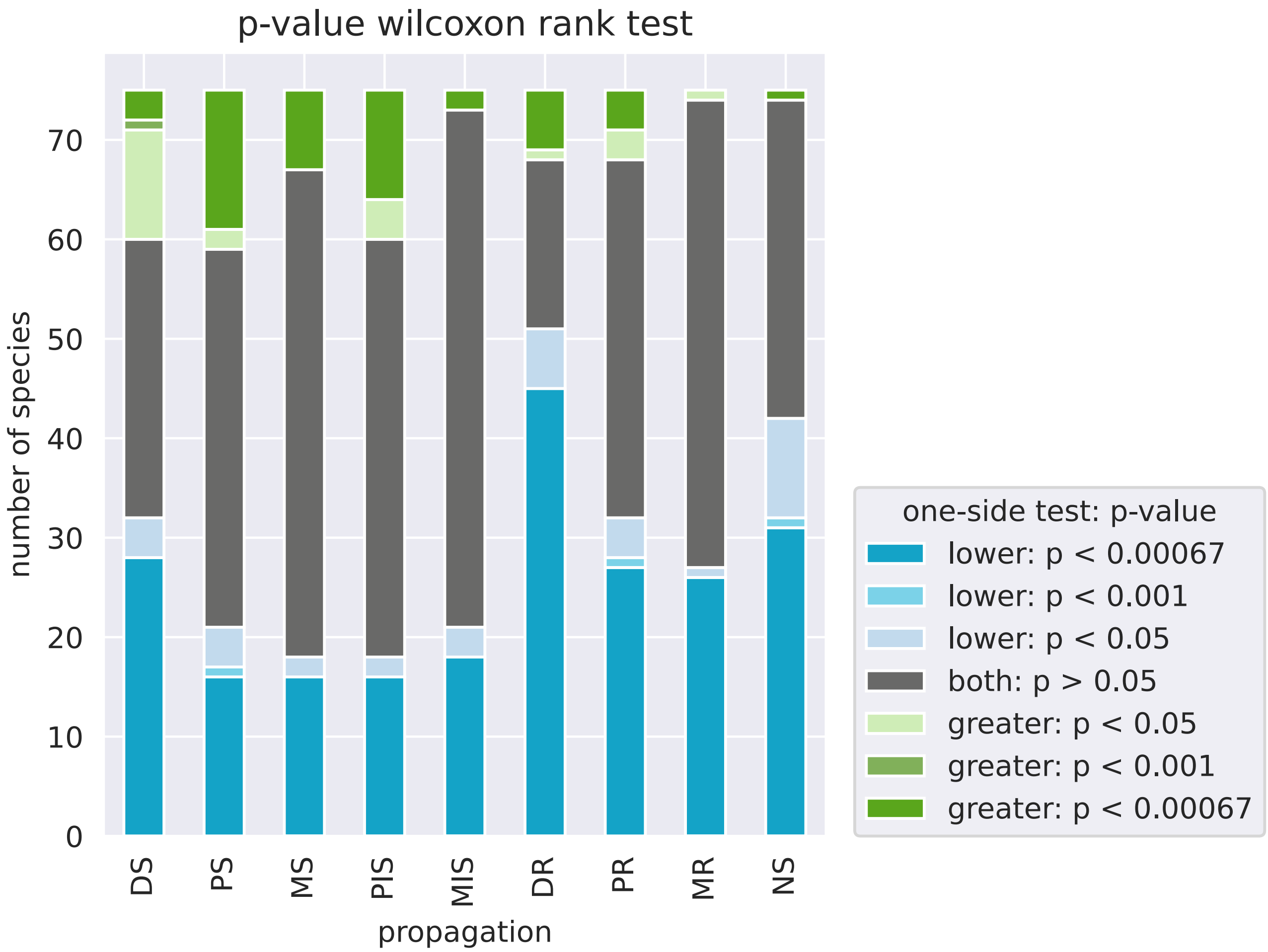


**Fig. S9.** Distribution of p-values from a one-sided Wilcoxon signed-rank test of whether the total investment $f_{l\cdot}$ of a species is larger/smaller in the last round where a species survived, than the investment of the ancestral species. There is one bar for each selection method, with 15 species x 5 sets of species for each bar. The alternative hypothesis is that difference in investment (ancestral-evolved) is greater (green) or less (blue) than zero. Data generated by the IBM. Data for DS is shown in Fig. 5.

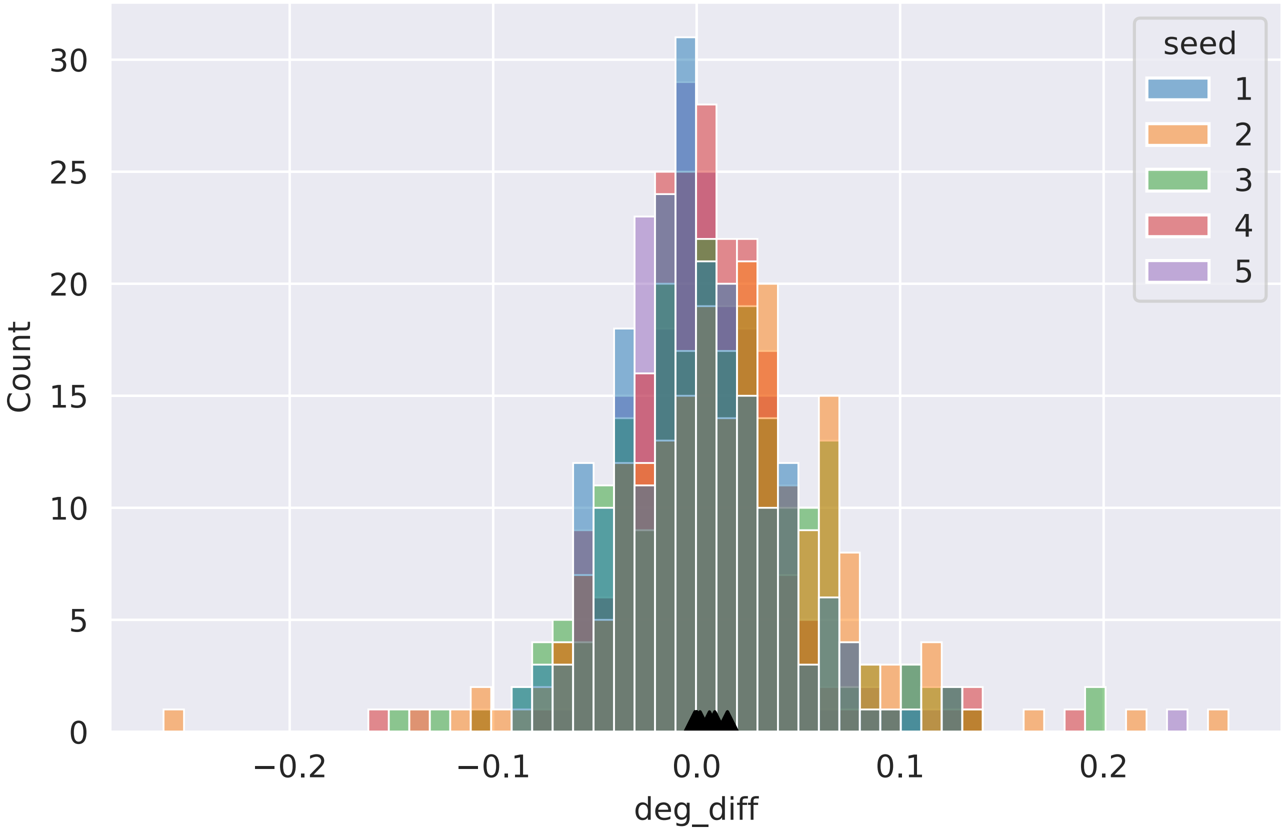


**Fig. S10.** Histogram of difference in max degradation between evolved and ancestral communities. Triangles indicate the mean values for each species set. Data generated by the IBM.

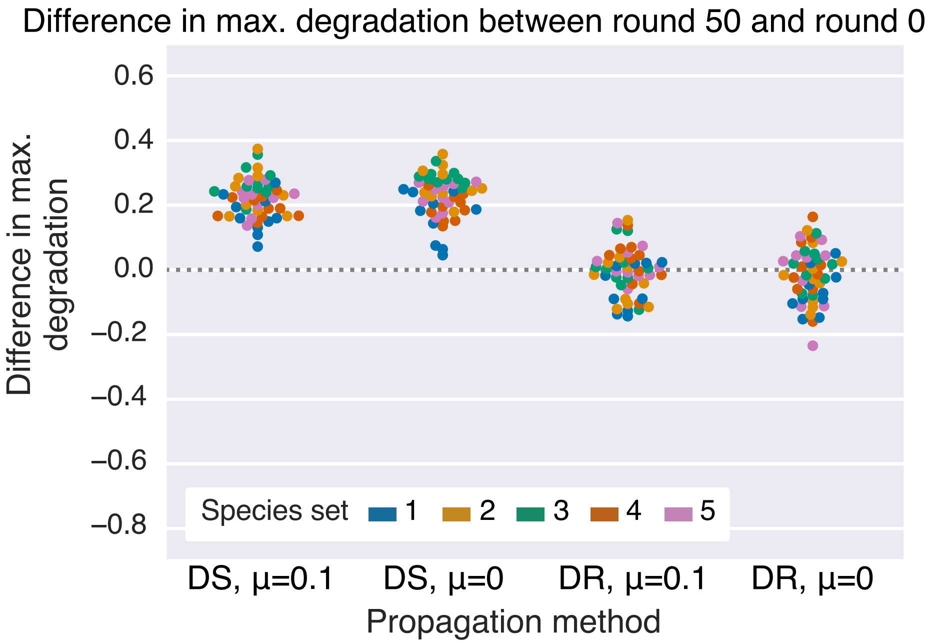


**Fig. S11.** The difference in maximum degradation between round 50 and round 0 for DS and DR, corresponding to Fig. 2, but varying the mutation rate from the default of $\mu = 0.1$ to $\mu = 0$. The performance of the method was not significantly affected by this change in mutation rate.

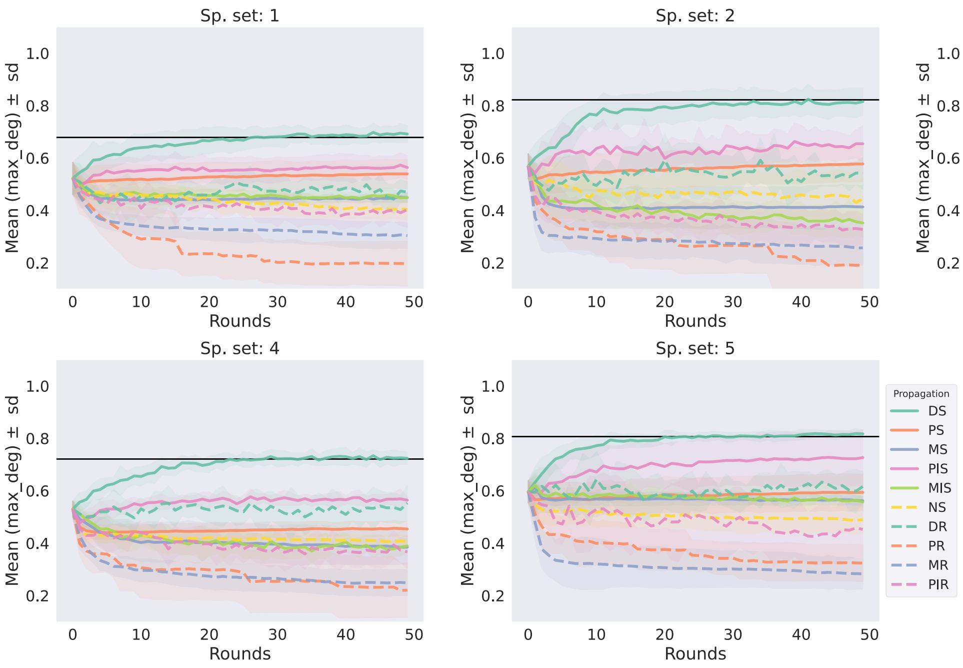

**Fig. S12.** Time-series of max. degradation over 50 rounds of selection, for the different propagation methods. Each plot corresponds to one species set and shows the maximum degradation in the meta-community averaged over repeats, with the standard deviation in shades of the corresponding color. For each species set, each repeat etc, the degradation score at transfer 50 forms the swarms in Fig. 2. The black line shows the degradation score of the best ancestral community out of the 32767 combinations. Data generated by the IBM.

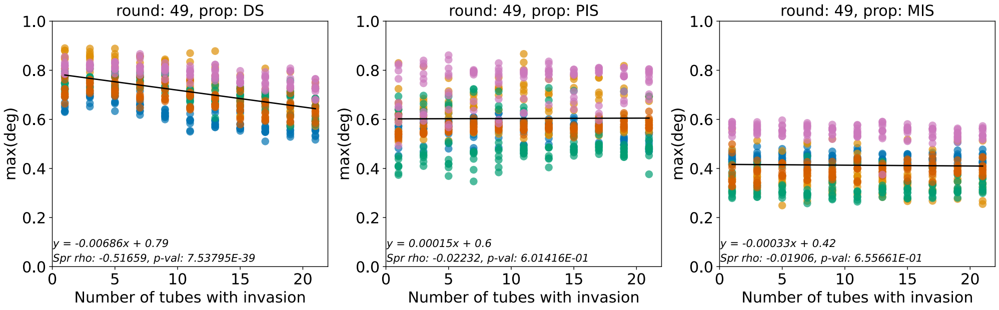

**Fig. S13.** Effect on max community degradation score from changing the number of communities to receive a migrating species. Data generated by the IBM.

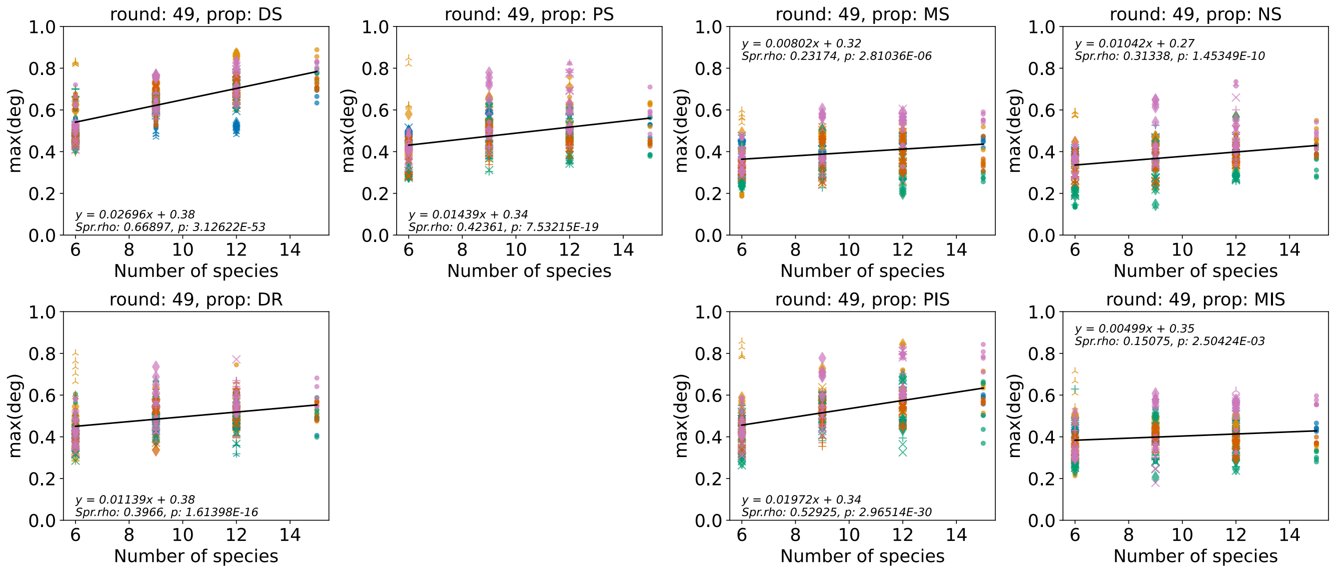


**Fig. S14.** Effect on max community degradation score from changing the number of species in the ancestral community. Different marker shape indicates sub-sample (1-5) for each species group of size 6, 9, 12. For 15 species we keep the original species sets. Data generated by the IBM.

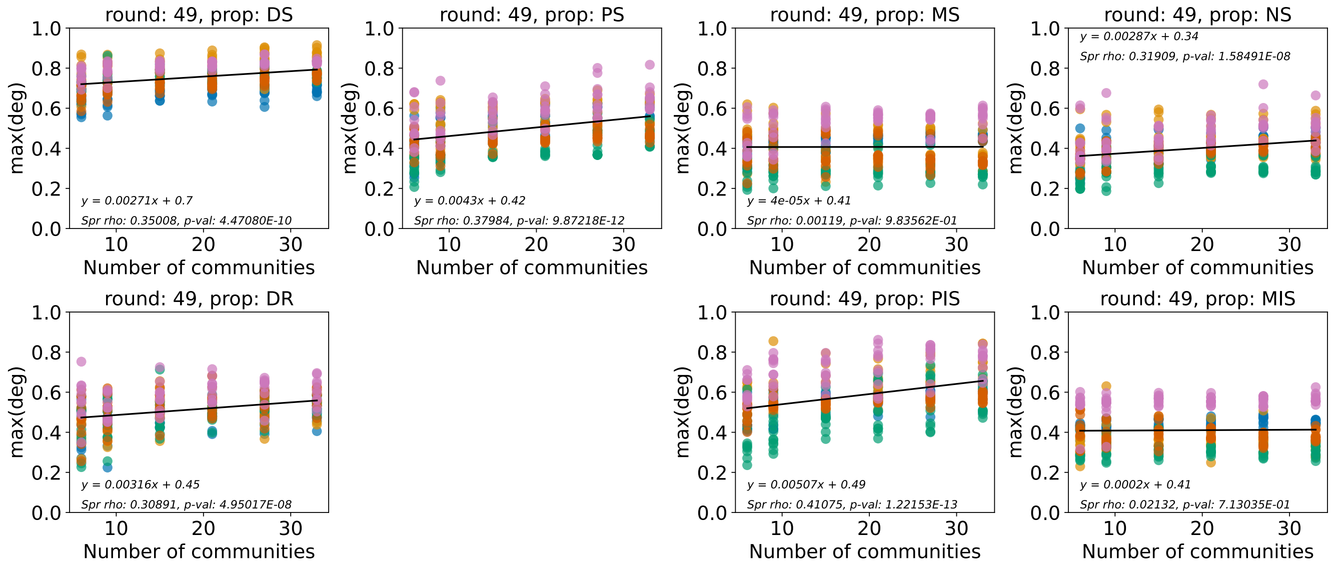


**Fig. S15.** Effect on max community degradation score from changing the number of communities. Data generated by the IBM.

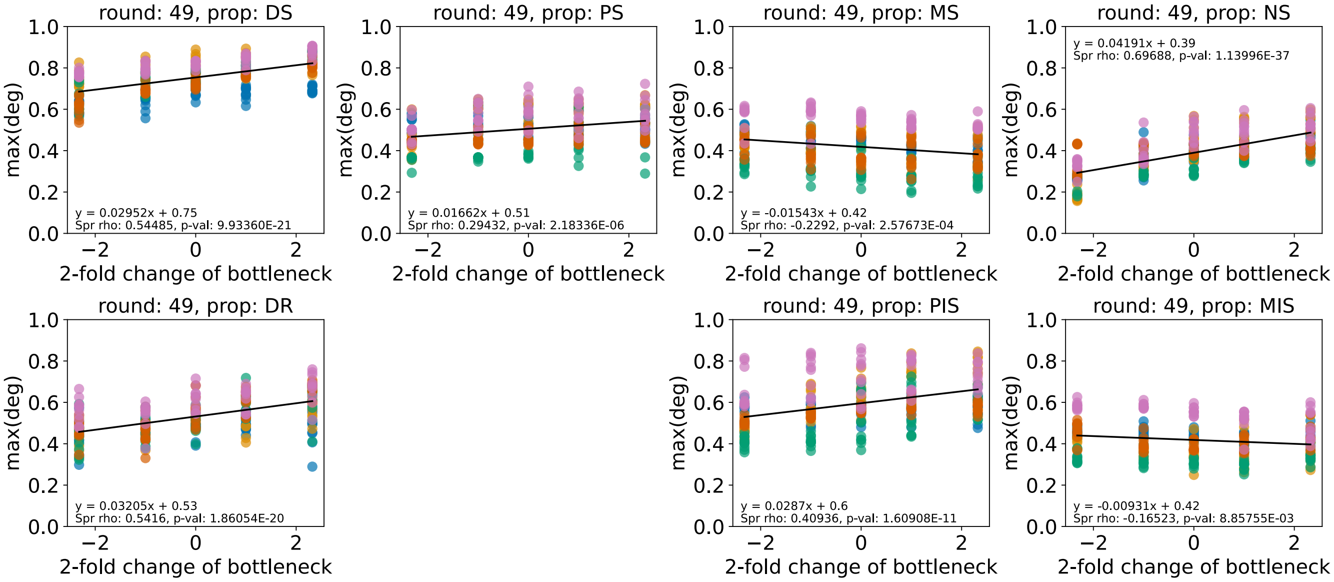


**Fig. S16.** Effect on max community degradation score from scaling the dilution factor or inoculum size. Data generated by the IBM.

#### Pseudo-code for implementation of models and selection methods

**Input:** A set of 15 species, defined by model parameters.

**Input:** Experimental parameters: Number of communities, time span $[t_0,\ t_{end}]$ of growth in batch. Community bottleneck $\beta = 1/3$, dilution ratio $d \in [0,\ 1]$. Initial conditions $S_i(t_0)$, $N_j(t_0)$, $T_k(t_0)$.

Assemble 21 communities by randomly drawing 4 species with replacement from the species set. Ensure that each species is present in at least one initial community.

**for** Each round of selection **do**

    // ***Population growth, interspecies competition and invasion of mutants***

    **for** Each community **do**

        Grow the communities for a time span $[t_0,\ t_{end}]$. (IBM implementation: Alg. 2, ODE implementation: Alg. 6).

        Save the population sizes $S_i(t_{end})$ for each strain $i$ in the community

        Save the end-state concentrations $T_k(t_{end})$ for each toxic compound $k$

        Compute the degradation score $D$ from $T_k(t_{end})$ by Eq. (4)

    **end**

    **// Propagate the communities by the chosen selection method**

    // Required parameters: the community bottleneck $\beta$, dilution ratio $d$

    // Required variables: degradation scores $D$ for each community

    *// For propagule method, follow Algs. 7 and 8*

    *// For migrant pool method, IBM only, follow Alg. 9*

    *// For disassembly method, follow Alg. 10*

    **// Replenish the substrates**

    Set $N_j(t_0) = N_0$ and $T_k(t_0) = T_0$ for all $j$, $k$

**end**

**Algorithm 1:** Overall flow of the selection simulations from population growth, community dynamics and mutations to propagation by the different selection methods.

**Input:** A community where each microbial strain $i$ is defined by the parameters in Tab. 1. Size of the inactive and active sub-populations $p_{i0}$, $p_{i1}$ and total population size $S_i = p_{i0} + p_{i1}$. Nutrient concentrations $N_j$, toxic compound concentrations $T_k$.
**Input:** Mutation parameters: mutation rate $\mu_{mut}$, trait deviation $\sigma_m$.

```
for Each time step do
    for Each community 1, ..., 21 do
        for Each strain i do
            // Maximum uptake of nutrients of this strain, used to scale the growth and degradation according to the
               concentration of nutrients
            max_uptake := Σ_j (n_ij if N_j > n_ij)
            // If some nutrients are depleted we re-scale n_ij to consume the remaining nutrients in higher amounts
            if N_j < n_ij then
                n_ij := 0
            end
            n̂_ij := n_ij / Σ_j (n_ij if N_j > n_ij)
            // The largest number of cells that can consume the scarcest nutrient at this time step, we just take its integer
               part
            S_i^max := int(min(N_j / n̂_ij if N_j > n̂_ij))
            // Toxic compound degradation
            if S_i^max > 0 then
                // Maximal population that can degrade
                P_i,tot := min(S_i^max, S_i)
                for Each compound T_k do
                    if T_k cannot be completely degraded in this time step then
                        T_k := T_k − P_i,tot · max_uptake · f_ik
                        for Each nutrient N_j do
                            N_j := N_j − n̂_ij · P_i,tot · f_ik
                        end
                    end
                    else
                        Degrade the remaining toxic compounds and consume the corresponding nutrients
                    end
                end
            end
            // Cell division step 1: Costly activation
            Alg. 3.
        end
    end
    // Cell division step 2: Replication
    Alg. 4.
    // Cell death
    Alg. 5.
end
```

**Algorithm 2:** Implementation of population growth, competition and mutations in the IBM model described in the section *Individual-based model*.

**Input:** Communities where each strain $i$ is defined by parameters in Tab. 1. Inactive and active sub-populations $p_{i0}$, $p_{i1}$. The maximal population $S_i^{max}$ that can afford to consume nutrients, based on their current availability. Re-scaled nutrient consumption rates $\hat{n}_{ij}$. Current nutrient concentrations $N_j$.

**Input:** Parameters: Initial nutrient concentration $N_0$.

*// Cell activation*

**if** $S_i^{max} > 0$ **then**

  *// Already activated cells consume nutrients*

  **if** $S_i^{max} \geq p_{i1}$ **then**

    $N_j := N_j - \hat{n}_{ij} \cdot p_{i1} \cdot (1 - \sum_k f_{ik})$

    $S_i^{max} := S_i^{max} - p_{i1}$

  **end**

  **else**

    Deactivate cells that cannot afford to stay activated, consume the corresponding nutrients for cells that remain activated, set $S_i^{max} := 0$

  **end**

  *// Newly activated cells*

  $cells\_activate :=$ Poisson$(a_i \cdot p_{i0} \cdot max_uptake \cdot (1 - \sum_k f_{ik}) \cdot \sum_j (\hat{n}_{ij} \cdot N_j / N_0))$

  **if** $cells\_activate > p_{i0}$ **then**

    $cells\_activate := p_{i0}$

  **end**

  **if** $cells\_activate > S_i^{max}$ **then**

    $cells\_activate := S_i^{max}$

  **end**

  $p_{i0} := p_{i0} - cells\_activate$

  $p_{i1} := p_{i1} + cells\_activate$

  **for** Each nutrient $N_j$ **do**

    $N_j := N_j - \hat{n}_{ij} \cdot cells\_activate \cdot (1 - \sum_k f_{ik})$

  **end**

**end**

**else**

  *Deactivate all the activated cells*

**end**

**return** *Populations* $p_{i0}$, $p_{i1}$ *for strain i, current nutrient concentrations* $N_j$

**Algorithm 3:** Activation of cells, the first step of cell division in the IBM described in Alg. 2.

**Input:** Communities where each strain $i$ is defined by parameters in Tab. 1. Inactive and active sub-populations $p_{i0}$, $p_{i1}$.

**for** Each community **do**
- **for** Each strain $i$ in the community **do**
  - // *Calculate the number of new cells appearing due to division*
  - $new\_cells$ :=Poisson $(p_{i1} \cdot r_i \cdot (1 - \sum_k f_{ik}))$
  - **if** $new\_cells > p_{i1}$ **then**
    - $new\_cells := p_{i1}$
  - **end**
  - // *Calculate how many* $new\_cells$ *will carry mutations*
  - $mutants$ :=Poisson $(new\_cells \cdot \mu_{mut})$
  - **if** $mutants > new\_cells$ **then**
    - $mutants := new\_cells$
  - **end**
  - $p_{i1} := p_{i1} - new\_cells$
  - $p_{i0} := p_{i0} + new\_cells \cdot 2 - mutants$
  - //*Mutation*
  - **for** Each new mutant **do**
    - Add a new strain $i$ to the community, with the model parameters of the ancestor and set $p_{i0} := 1$ and $p_{i1} := 0$
    - Decide which $f_{ik}$ to mutate by drawing from Bernoulli$\left(\frac{1}{N_{tox}}\right)$ for each $f_{ik}$; ensure that at least one $f_{ik}$ mutates
    - **for** Each successful draw **do**
      - Multiply the chosen $f_{ik}$, by a factor $x \sim lognormal(0.0, \sigma_m)$
      - Re-scale so $\sum_k f_{ik} \leq 1$, if needed
    - **end**
  - **end**
- **end**

**end**

**return** *Populations* $p_{i0}$, $p_{i1}$ *for each strain i, including the new ones resulted from mutation.*

**Algorithm 4:** Replication and mutation, second step of cell division of the IBM described in Alg. 2.

**Input:** Communities where each strain $i$ is defined by parameters in Tab. 1. Inactive and active sub-populations $p_{i0}$, $p_{i1}$. Current concentrations $T_k$ of toxic compounds. Death rates $m_{ik}$ and $K$ constant for the Hill function.

**for** Each community **do**
- **for** Each strain $i$, looping over the community in reverse order **do**
  - $p_{i0} := p_{i0} -$Poisson$\left(p_{i0} \cdot \sum_k (m_{ik} \cdot \frac{T_k^2}{T_k^2 + K^2})\right)$; ensure that $p_{i_0} \geq 0$
  - $p_{i1} := p_{i1} -$ Poisson $\left(p_{i1} \cdot \sum_k (m_{ik} \cdot \frac{T_k^2}{T_k^2 + K^2})\right)$; ensure that $p_{i_1} \geq 0$
  - **if** $p_{i0} + p_{i1} = 0$ **then**
    - remove strain $i$ from the community
  - **end**
- **end**
- Randomly shuffle strains in community

**end**

**return** *Populations* $p_{i0}$, $p_{i1}$ *for each strain* $i$.

**Algorithm 5:** Cell death of the IBM described in Alg. 2.

**Input:** Strains with model parameters from S4 for each population $i$.
**Input:** Mutation parameters: rate $\mu_{mut}$, trait deviation $\sigma_m$.
**Input:** Experimental parameters: Number of toxic compounds $N_{tox}$, initial concentrations $N_0$, $T_0$ of nutrients and toxic compounds, initial population size $S_0$. Time span for growth $[t_0,\ t_{end}]$
**for** Each community **do**
  *// Growth and competition within one round*
  Solve the equations Eq. (S1)–Eq. (S3) for a time span $[t_0,\ t_{end}]$.
  Save the end states $S_i(t_{end})$, $N_j(t_{end})$, $T_k(t_{end})$.
  // Mutations, ODE model
  **for** Each strain $i$, with probability $\mu_{mut}$ **do**
    Copy the species parameters to an empty place in the list of populations
    Choose a $f_{ik}$ at random by drawing from Bernoulli$(1/N_{tox})$ for each $k = 1,\ \ldots,\ 10$
    For each chosen $f_{ik}$, multiply by a factor $x_k \sim lognormal(0.0,\ \sigma_m)$
    Set the inoculum size to $S_0$
  **end**
**end**
**return** $S_i(t_{end})$, $N_j(t_{end})$, $T_k(t_{end})$, $f_{ik}$

**Algorithm 6:** Implementation of population growth, competition and mutations in the ODE model described in the section *Population-level model*. To solve the equations, we use *dopri5* from the SciPy library (42, 45).

**Input:** Communities with populations $S_i$ and degradation scores $D$. End states $T_k(t_{end})$.
**Input:** Experimental parameters: selection bottleneck $\beta = 1/3$, dilution ratio $d$.
Rank the communities by degradation $D$
Select the top $N_\beta = 7$ of communities with the highest ranks.
*// Re-populate the new set of tubes*
Allocate $1/\beta$ new tubes for each selected community
**for** Each selected community $1,\ 2,\ \ldots,\ 7$ **do**
  **for** Each population $S_i$ in the selected community **do**
    *// Dilute the population*
    Dilute $S_i(t_0) := d \cdot S_i(t_{end})$
    **if** $S_i(t_0) < 1.0$ **then**
      *// The population is extinct*
      Set $S_i(t_0) := 0.0$
      Remove all species parameters from the community
    **end**
    Copy model parameters and population sizes $S_i(t_0)$ of each strain in the parent communities to each of the $1/\beta$ offspring communities
  **end**
**end**

**Algorithm 7:** Implementation of the propagule selection method for the ODE model.

**Input:** Communities with degradation scores $D$ and strains $S_i$ with total population $S_i = p_{i0} + p_{i1}$. End-state concentration of toxic compounds $T_k(t_{end})$.
**Input:** Experimental parameters: selection bottleneck $\beta = 1/3$, dilution ratio $d$.

```
Rank the communities by D
Select the top N_β = 7 of communities with the highest ranks
// Re-populate the new set of tubes
Allocate 1/β new tubes for each selected community
for Each selected community 1, 2, ..., 7 do
    for Each strain i in the community do
        // Deactivate cells
        p_i0(t_end) = S_i(t_end)
        p_i1(t_end) = 0
        // Dilute the population, new cells will be inactivated
        p_i0(t_0) :=Poisson(d · S_i(t_end))
        if p_i0(t_0) > S_i(t_end) then
            p_i0(t_0) = S_i(t_end)
        end
        if p_i0(t_0) > 0 then
            Append strain i with population p_0(t_0) to the new tube
        end
        // Delete selected cells to not chose them again
        S_i(t_end) = S_i(t_end) − p_i0(t_0)
    end
end
```

**Algorithm 8:** Implementation of the propagule selection method for the IBM.

**Input:** Communities with degradation scores $D$ and strains $S_i$ with total population $S_i = p_{i0} + p_{i1}$. End-state concentration of toxic compounds $T_k(t_{end})$.

**Input:** Experimental parameters: selection bottleneck $\beta = 1/3$, dilution ratio $d$.

Rank the communities by $D$

Select the top $N_\beta = 7$ communities with the highest ranks, and pool their populations.

*// Re-populate the new set of tubes*

Allocate 21 new tubes

**for** Each offspring community $1, 2, \ldots, 21$ **do**

  **for** Each strain $i$ in the pool **do**

    *// Deactivate cells*

    $p_{i0}(t_{end}) = S_i(t_{end})$

    $p_{i1}(t_{end}) = 0$

    *// Dilute the population, new cells will be inactivated*

    Draw $p_{i0}(t_0)$ from Poisson $(\frac{d}{N_\beta} \cdot S_i(t_{end}))$

    **if** $p_{i0}(t_0) > S_i(t_{end})$ **then**

      $p_{i0}(t_0) = S_i(t_{end})$

    **end**

    **if** $p_{i0}(t_0) > 0$ **then**

      Append strain $i$ with population $p_{i0}(t_0)$ to the new tube

    **end**

    *// Delete selected cells to not chose them again*

    $S_i(t_{end}) = S_i(t_{end}) - p_{i0}(t_0)$

  **end**

**end**

**Algorithm 9:** Implementation of the migrant pool selection method for the IBM

**Input:** Communities with populations $S_i$ (in the IBM $S_i = p_{i0} + p_{i1}$) with model parameters from (Tab. 1, Tab. S4) and degradation scores $D$.

**Input:** Experimental parameters: selection bottleneck $\beta = 1/3$, initial population size $S_0$ and number of new communities to emigrate species from $N_{emi} = 5$, and immigrate species to $N_{immi} = 5$.

// *Rank the communities*

Rank the communities by the degradation score $D$

// *Update the fossil record with the top communities in this round*

**for** Each selected community $1, 2, \ldots, 7$ **do**

  **for** Each species $l$ in the selected community **do**

    **if** The record of species $l$ is not yet updated in this round of selection **then**

      Add all strains $i$ of species $l$ to the fossil record, including the corresponding parameters and population sizes

    **end**

  **end**

**end**

// *Propose new communities in proportion to their degradation scores and survival*

Follow Alg. 11

// *Emigration*

Draw $N_{emi} = 5$ communities with uniform probability

**for** Each chosen community $1, \ldots, N_{emi}$ **do**

  // *Find emigrating species*

  $Number\_of\_emigrants = 1 + \text{Poisson}(0.5)$

  Verify that at least one species will remain

  **for** Each emigrant **do**

    Choose the emigrant at random, with priority for species that occur in more than one community

    Remove all strains of this species from the community

  **end**

**end**

// *Immigration*

Draw $N_{immi} = 5$ communities with uniform probability

**for** Each chosen community $1, \ldots, N_{immi}$ **do**

  // *Find immigrating species*

  $Number\_of\_immigrants = 1 + \text{Poisson}(0.5)$

  **for** Each immigrant **do**

    **if** *There are species that do not feature in any community* **then**

      Choose one of them at random

    **end**

    **else**

      Choose a species that is not already in the community with uniform probability

    **end**

    Take all strains of the species from the species record, add them to the offspring community

    Set the population size to $S_0$ (approximately $S_0$ in the IBM, see Alg. 11), in proportion to strain relative abundance

  **end**

**end**

**Algorithm 10:** Implementation of the disassembly method.

**Input:** The subset of selected communities, their degradation scores $D$. Population sizes $S_i$ of all strains (in the IBM, $S_i = p_{i0} + p_{i1}$).

**Input:** Parameters: Number of species $N_{spc}$ that were inoculated in the different communities. Inoculum size per species $S_0$.

// Scale degradation scores by species extinctions

**for** Each selected community $1, 2, \ldots, 7$ **do**
    // Count the number of surviving species in this community
    Set the extinction counter $E = 0$
    **for** Each species $l$ in the community **do**
        **if** $S_i = 0$ *for all strains of species* $l$ **then**
            Set $E := E + 1$
            *// Re-introduce species* $l$ *from the record*
            Take all strains of species $l$ from the most recent record
        **end**
    **end**
    // Scale the degradation score $D$ by the fraction of surviving species
    Set $\hat{D} := D \cdot (N_{spc} - E)/N_{spc}$
**end**

*// Calculate a probability distribution based on degradation scores*

Set $p_n := \hat{D}_n / \sum_m \hat{D}_m$ for each community $n$

*// Propose new communities randomly in proportion to* $p_n$

**for** Each offspring community **do**
    Choose a parental community at random, by the probability distribution $p_n$
    **for** Each species $l$ in the parental community **do**
        Copy the growth parameters and $f_{ik}$ of all strains $i$ to the offspring community
        // ODE: Set the population size of each species to $S_0$ in total, in proportion to the relative abundance of the strains.
        // IBM: Sample approximately $S_0$ cells with replacement as follows:
        **for** Each strain $i$ (with population $S_i$) of species $l$ (with population $S_l$) **do**
            *//Deactivate cells*
            $p_{i0}(t_{end}) = S_i(t_{end})$
            $p_{i1}(t_{end}) = 0$
            *// Draw new population*
            Draw $p_{i0}(t_0)$ from Poisson $(S_0 \cdot \frac{S_i}{S_l})$
            **if** $p_{i0}(t_0) > 0$ **then**
                Add strain $i$ with population $p_{i0}(t_0)$ to the new community
            **end**
        **end**
    **end**
**end**

**Algorithm 11:** Method to propose new communities based on their degradation scores from the previous round. Called by Alg. 10

| Parameter | Description | Sampled from |
|---|---|---|
| $l_i$ | Species ID of strain $i$ | |
| $r_{ij}$ | Maximum growth rate with respect to nutrient $j$ | $\text{Uni}(0.01,\ 0.1)$ Sparse |
| $K_N$ | Half-saturation constant for nutrients | $K_N = 10$ (fixed) |
| $m_{ik}$ | Maximum death rate with respect to toxic compound $k$ | $\text{Uni}(10^{-4},\ 10^{-3})$ Sparse |
| $K_T$ | Half-saturation constant for toxic compounds | $K_T = 10$ (fixed) |
| $f_{ik}$ | Fraction of amassed nutrients that are invested into degradation of toxic compound $k$ | $\text{Uni}(0,1)$, Sparse Rescaled so that $\sum_k f_{ik} = \text{Uni}(0,1)$ |
| $Y_i$ | (Average) biomass yield with respect to the nutrients | $\text{lognormal}(\log(10^{-3}), \log(5))$ |
| $\delta_i$ | (Average) degradation efficiency with respect to the toxic compounds | $\text{lognormal}(\log(10^{-4}), \log(5))$ |

**Table S4.** Parameters defining a microbial strain $i$ in the ODE model. Growth and degradation parameters in relation to nutrients and toxic compounds $N_j$ and $T_k$. All parameters are assumed to be positive, and the investment $f_{ik}$ is limited to the interval $[0,\ 1]$. The matrices of growth rates, death rates and degradation investment $r_{ij}$, $m_{ik}$ and $f_{ik}$ are made sparse by multiplying them by matrices drawn from Bernoulli$(0.5)$, i.e. flipping a coin for each entry. In this way, each species takes up approximately half of the nutrients, is affected by half of the toxic compounds and degrades half of the toxic compounds.

### Supplementary methods

**ODE model.** As in the IBM, the ODE model simulates well-mixed batch cultures with nutrients and toxic compounds, extending a previous model (29). In the model, the population size $S_i$ of each strain $i$ in a community grows in relation to the concentrations of nutrients $N_j$ and decline by the toxic compounds $T_k$ (Fig. S1) by the model parameters in Table S4. Growth, death, nutrient uptake and degradation is described by the following ODE system:

$$\frac{dS_i}{dt} = \left( (1 - \sum_k f_{ik}) \rho_i(\mathbf{N}) - \mu_i(\mathbf{T}) \right) S_i \quad \textbf{(S1)}$$

$$\frac{dN_j}{dt} = -\sum_i \frac{\rho_i(N_j)}{Y_i} S_i \quad \textbf{(S2)}$$

$$\frac{dT_k}{dt} = -T_k \sum_i f_{ik} \delta_i \rho_i(\mathbf{N}) S_i \quad \textbf{(S3)}$$

The bold-face $\mathbf{N}$, $\mathbf{T}$ denote the vectors of all nutrients and toxic compounds, respectively. We assume Monod and Hill functions for the per-capita growth and death rates $\rho_i$, $\mu_i$.

$$\rho_i(\mathbf{N}) = \sum_j r_{ij} \frac{N_j}{N_j + K_N} \quad \textbf{(S4)}$$

$$\mu_i(\mathbf{T}) = \sum_k m_{ik} \frac{T_k^2}{T_k^2 + K_T^2} \quad \textbf{(S5)}$$

The system of equations Eq. (S1)–Eq. (S3) is solved with a standard ODE solver (`dopri5`, (42, 45)) for 100 time steps with initial conditions $S_i(t_0) = 100$, $N_j(t_0) = 100$ and $T_k(t_0) = 100$ for all $i$, $j$, $k$.
The investment $f_{ik}$ can mutate to form different strains of the same species. When this happens, we add a new population equation of the type Eq. (S1) to the ODE system, with the same parameters $r_{ij}$, $m_{ik}$, $Y_i$ and $d_i$ as the ancestor but with the modified $f_{ik}$. To not make the system of equations too large, we have limited the number of strains to 28 per community. We estimate this to be enough since we expect mutants to rapidly replace their ancestral strains if their growth rate is higher, and

otherwise disappear rapidly. If there are already 28 strains in a community, then no more mutants are allowed. Otherwise, when communities are propagated to the next round of growth, any surviving strain can have a mutant with probability $0.05$. Having chosen which strains $i$ to mutate, we pick one or more traits $f_{ik}$ at random and multiply them by numbers drawn at random from lognormal$(0,\ 0.4)$ and ensure that both the mutated traits $f_{ik}$ and the total investment $f_{i\cdot}$ falls in the $[0,\ 1]$ interval. The mutant receives the same $r_{ij}$, $m_{ik}$, $Y_i$ and $d_i$ parameters as its ancestor and is introduced with population size $100$, the same as the initial population before the first round of growth. This population size is chosen relatively high, in order to speed up the competition between ancestor and mutant strain.
